# Theta and Gamma Bands Encode Acoustic Dynamics over Wide-ranging Timescales

**DOI:** 10.1101/547125

**Authors:** Xiangbin Teng, David Poeppel

## Abstract

Natural sounds have broadband modulation spectra and contain acoustic dynamics ranging from tens to hundreds of milliseconds. How does the human auditory system encode acoustic information over wide-ranging timescales to achieve sound recognition? Previous work (Teng et al., 2017) demonstrated a temporal coding preference in the auditory system for the theta (4 – 7 Hz) and gamma (30 – 45 Hz) ranges, but it remains unclear how acoustic dynamics between these two ranges is encoded. Here we generated artificial sounds with temporal structures over timescales from ~200 ms to ~30 ms and investigated temporal coding on different timescales in the human auditory cortex. Participants discriminated sounds with temporal structures at different timescales while undergoing magnetoencephalography (MEG) recording. The data show robust neural entrainment in the theta and the gamma bands, but not in the alpha and beta bands. Classification analyses as well as stimulus reconstruction reveal that the acoustic information of all timescales can be differentiated through the theta and gamma bands, but the acoustic dynamics in the theta and gamma ranges are preferentially encoded. We replicate earlier findings of multi-time scale processing and further demonstrate that the theta and gamma bands show generality of temporal coding across all timescales with comparable capacity. The results support the hypothesis that the human auditory cortex primarily encodes auditory information employing neural processes within two discrete temporal regimes.

**Significance:** Natural sounds contain rich acoustic dynamics over wide-ranging timescales, but perceptually relevant regularities often occupy specific temporal ranges. For instance, speech carries phonemic information on a shorter timescale than syllabic information at ~ 200 ms. How does the brain efficiently ‘sample’ continuous acoustic input to perceive temporally structured sounds? We presented sounds with temporal structures at different timescales and measured cortical entrainment using magnetoencephalography. We found, unexpectedly, that the human auditory system preserves high temporal coding precision on two non-overlapping timescales, the slower (theta) and faster (gamma) bands, to track acoustic dynamics over all timescales. The results suggest that the acoustic environment which we experience as seamless and continuous is segregated by discontinuous neural processing, or ‘sampled.’

## Introduction

Natural sounds contain rich acoustic dynamics over a wide temporal range reflected in their broadband modulation spectra (Nelken et al., 1999; Lewicki, 2002; Singh and Theunissen, 2003; Narayan et al., 2006), whereas perceptually relevant information embedded in such wide-ranging acoustic dynamics often occupies specific timescales. For example, syllabic information in speech unfolds over ~200 ms temporal windows while phonemic information is conveyed in ~50 ms (Rosen, 1992). How, then, does the auditory system efficiently extract relevant information to achieve sound recognition? One strategy would be to process acoustic information at every timescale equally and derive the appropriate perceptual representations by integrating information across all timescales without preferring any particular timescales. A different strategy would be to selectively analyze slowly varying auditory attributes, carried over a longer timescale, to guarantee sufficient information for perceptual analysis, and concurrently to extract fast changing dynamics on a shorter timescale to preserve temporal resolution (Poeppel, 2003; Boemio et al., 2005; Giraud and Poeppel, 2012; Teng et al., 2017). To adjudicate between these alternatives, we test how the auditory system is entrained by acoustic dynamics over wide-ranging timescales.

Neural entrainment reflects the dynamics of neural populations at different timescales evoked by sensory stimuli and is thought to reveal the corresponding temporal characteristics of sensory processing (VanRullen, 2006; Henry and Obleser, 2012; Henry et al., 2014; VanRullen et al., 2014). Previous studies used various stimuli, such as speech, music, and amplitude- or frequency-modulated sounds, and found robust neural entrainment in auditory cortical areas below 10 Hz, suggesting a high temporal coding capacity in the low frequency range (Luo and Poeppel, 2007; Lakatos et al., 2008; Kerlin et al., 2010; Besle et al., 2011; Cogan and Poeppel, 2011; Ding and Simon, 2012; Henry and Obleser, 2012; Kayser et al., 2012; Ng et al., 2012; Wang et al., 2012; Ding and Simon, 2013; Herrmann et al., 2013; Lakatos et al., 2013; Peelle et al., 2013; Doelling et al., 2014; Henry et al., 2014; Kayser et al., 2015; Riecke et al., 2015; Zoefel and VanRullen, 2015). On the other hand, there is evidence suggesting that the low gamma band plays an important role in syllable processing and comprehension of speech (Palva et al., 2002; Shahin et al., 2009; Kerlin et al., 2010; Peña and Melloni, 2011; Morillon et al., 2012; Gross et al., 2013). Previous experiments examined neural entrainment at the low and high frequency ranges and demonstrated that both the neural theta and gamma bands, but not the alpha band, robustly encode acoustic information (Luo and Poeppel, 2012; Teng et al., 2017). However, two key mechanistic questions remain unresolved: how are acoustic dynamics between the theta and gamma ranges encoded? Is acoustic information encoded preferentially with higher precision in the theta band range, given the higher magnitude of neural responses in the low frequency range? Or is temporal coding capacity comparable between the theta and gamma bands?

Here, we test the temporal coding capacity of the human auditory cortex from 4 to 45 Hz by measuring neural entrainment. Building on earlier work (Teng et al., 2017), we generated acoustic stimuli with modulation rates centered at theta (4 - 7 Hz), alpha (8 – 12 Hz), beta1 (13 – 20 Hz), beta2 (21 – 30 Hz) and low gamma bands (31 - 45) and conducted a match-to-sample task to evaluate listeners’ discriminability of different modulation rates. Classification and decoding analyses on the MEG data were performed to test which neural frequency bands preserve high temporal coding capacity and faithfully encode acoustic dynamics of different timescales. We show that the theta and gamma bands encode acoustic information with comparably high precision, while the alpha and beta bands manifest limited temporal resolution. Next, the stimuli modulated at all the timescales can be classified using the theta and gamma bands. Finally, the source localization results show that the neural activity of both the theta and gamma band originate from similar auditory cortical areas.

## Methods

### Ethics statement

The study was approved by the New York University Institutional Review Board (IRB# 10–7277) and conducted in conformity with the 45 Code of Federal Regulations (CFR) part 46 and the principles of the Belmont Report.

### Participants

Sixteen right-handed participants (9 females; age range: 23 - 41) took part in the experiment. Handedness was determined using the Edinburgh Handedness Inventory (Oldfield, 1971). All participants had normal hearing and reported no neurological deficits. We excluded the data from one participant because of noise issues during neurophysiological recording; therefore, the analysis included the data from 15 participants (9 females; age range: 23 - 35).

### Stimuli

We generated five types of stimuli building on methods used in previous studies (Boemio et al., 2005; Luo and Poeppel, 2012; Teng et al., 2017). Each stimulus was 2 s long and generated by concatenating narrow-band frequency-modulated segments, each of which consisted of 100 sinusoids with randomized amplitude, phase, and frequency. The bandwidth of segments was 100 Hz (within a critical band at the center frequencies used). The to-be-concatenated segments for each stimulus type were drawn from a Gaussian distribution with means of 190, 100, 62, 41 and 27 ms, and with standard deviations of 30, 15, 6, 4 and 3 ms, respectively. Hence the distribution of the segment durations aligned with the range of periods of theta (4-7 Hz), alpha (8-12 Hz), beta1 (13 – 20 Hz), beta2 (20 – 30 Hz), and low gamma (30-45 Hz) neural bands (Fig. 1*A*). The frequency-modulated segments could sweep up from 1000 Hz to 1500 Hz or down from 1500 Hz to 1000 Hz. To be concise, hereafter we refer to the stimulus type with mean segment duration of 190 ms as a “ theta (*θ*) sound”, the stimulus type with mean segment duration of 100 ms as an “alpha (*α*) sound”, the stimulus type with mean segment duration of 62 ms as a “beta1 (*β1*) sound”, the stimulus type with mean segment duration of 41 ms as a “beta2 (*β2*) sound” and the stimulus type with mean segment duration of 27 ms as a “gamma (*γ*) sound.”

**Figure 1.**
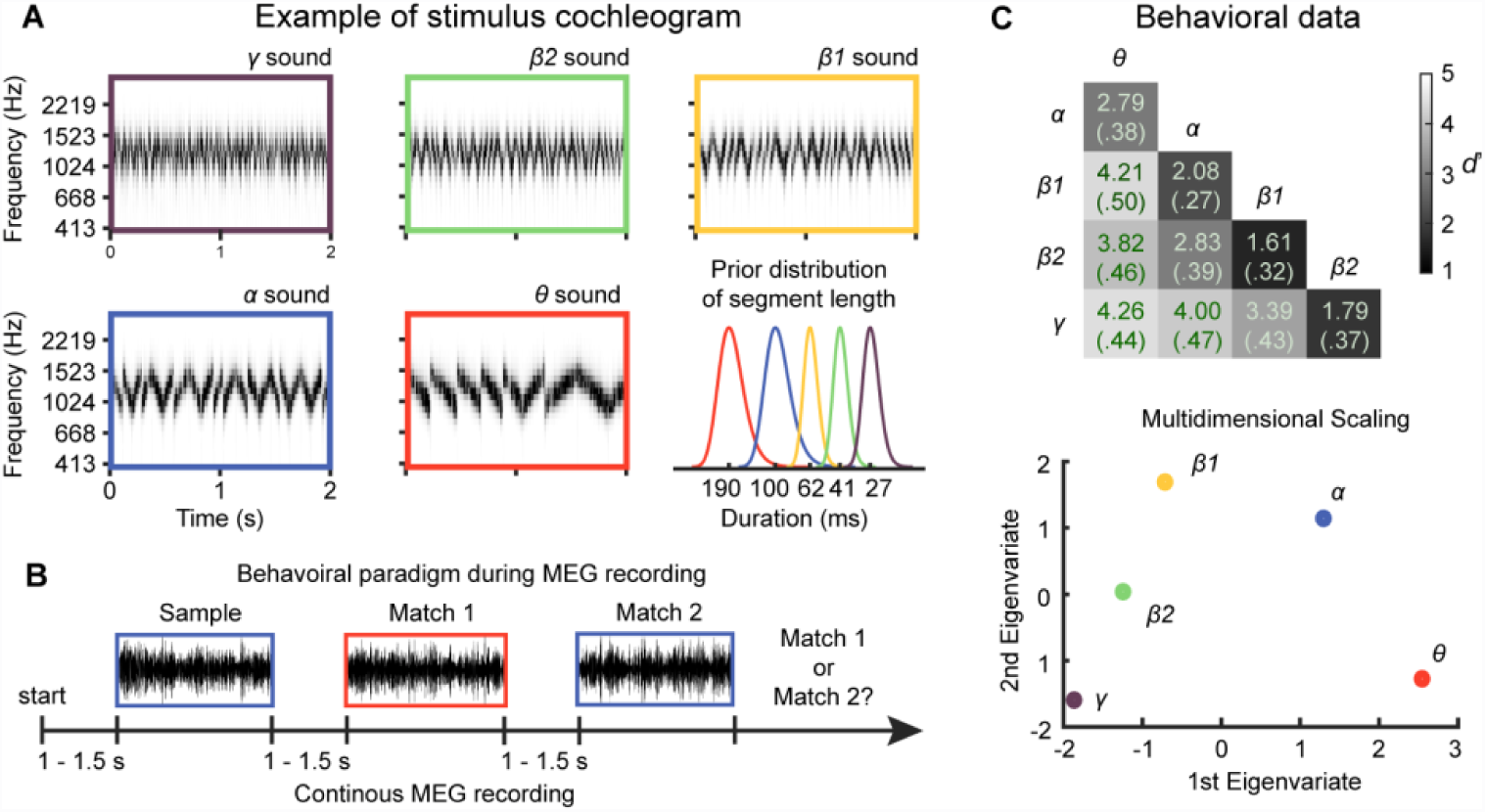
Experimental procedure and behavioral results. ***A***, Cochleograms of five stimulus types. The cochleogram of an example frozen sound from each stimulus type is shown from upper left to bottom middle: *θ* sound (modulation rate of 4 - 7 Hz), *α* sound (8 - 12 Hz), *β1* sound (13 – 20 Hz), *β2* sound (21 – 30 Hz) and *γ* sound (31 – 45 Hz). Their prior distributions of segment durations are shown in the bottom right panel. The color scheme codes for each stimulus type and is used consistently in the following figures. ***B***, Behavioral paradigm during MEG recording. Participants performed a match-to-sample task to differentiate different stimulus types (modulation rates). The distinct sounds were presented only in the sample interval and the frozen sounds in the match intervals. ***C***, Behavioral results. Upper panel shows group-averaged d-prime values in a form of confusion matrix of different pairs of stimulus types. The gray scale codes for d-prime values; the number in each cell represents group mean and the standard error of mean across participants (in parentheses). The results of multidimensional scaling on the group-averaged d-prime values are plotted in the lower panel and illustrate perceptual distance between different stimulus types. Data points represent each stimulus type.

For each stimulus type, we generated three samples with different modulation phases. For example, for the stimulus type with a modulation rate in the theta band, the *θ* sound, we generated three *θ* sounds with the same modulation rate but different modulation phases. The cochleogram of one example of the each stimulus type (Ellis, 2009) and the corresponding prior distribution of segment duration for each stimulus type are illustrated in Fig. 1*A*. Therefore, we have three sounds for each stimulus type and 15 sounds in total for five stimulus types. The 15 sounds were presented repeatedly in the experiment and named ‘frozen’ sounds. In addition, we generated 40 sounds with distinct modulation phases for each stimulus type. Each of the 40 sounds was presented only once and therefore named ‘distinct’ sounds, to indicate that each sound for one stimulus type has different modulation phases from the other sounds. In total, we have 200 distinct sounds for five stimulus types.

During MEG recording, participants performed a match-to-sample task to differentiate stimulus types (modulation rates), as illustrated in Fig. 1*B*. On each trial, participants were first required to focus on a white fixation cross in the center of a black screen. Then the screen showed a word in yellow, ‘sample’, and a sample stimulus was presented simultaneously, which was a distinct sound from one of the five stimulus types. After the sample stimulus was over, the screen showed the word ‘match’ and a pair of ‘match’ stimuli selected from the frozen sounds was presented, one of which matched the modulation rate of the sample stimulus. After the second match stimulus was presented, the participants were required to choose by pressing one of two buttons which interval in the match pair matched the sample stimulus. After the response, the next trial started in 1 ~ 1.5 s. The intervals between all three stimuli (sample stimulus and two match stimuli) were uniformly distributed between 1 to 1.5 s.

During the match-to-sample task, in the sample intervals, we presented 40 distinct sounds of each stimulus type; in the match intervals, two frozen sounds of each stimulus type were presented 27 times and one frozen sound 26 times. For each pair of stimulus types in comparison, 40 trials were presented to test listeners’ discriminability, with 20 trials having one stimulus type of the pair as the sample stimulus and 20 trials having the other stimulus type as the sample stimulus. In total, 200 trials were presented, which contained 200 (5 stimulus types * 40) distinct sounds as the sample stimuli and 400 (5 stimulus types * (27 + 27 + 26)) frozen sounds as the match stimuli.

All the stimuli were normalized to ~65 dB SPL and delivered through plastic air tubes connected to foam ear pieces (E-A-R Tone Gold 3A Insert earphones, Aearo Technologies Auditory Systems).

### MEG recording and channel selection

MEG signals were measured with participants in a supine position and in a magnetically shielded room using a 157-channel whole-head axial gradiometer system (KIT, Kanazawa Institute of Technology, Japan). A sampling rate of 1000 Hz was used with an online 1-200 Hz analog band-pass filter and a notch filter centered around 60 Hz. After the main experiment, participants were presented with 1 kHz tone beeps of 50 ms duration as a localizer to determine their M100 evoked responses, which is a canonical auditory response (Roberts et al., 2000). Twenty channels with the largest M100 response of both hemispheres (10 channels in each hemisphere) were selected as auditory channels for each participant individually.

### Behavioral data analysis

Behavioral data analysis was conducted in MATLAB using the Palamedes toolbox 1.5.1 (Prins and Kingdom, 2009). For each pair of stimulus types, we averaged correct responses and then converted the percentage correct to *d*′ assuming an independent observer model and an unbiased observer. To avoid infinite d-prime values, a half artificial incorrect trial was added in the case that all trials were correct; a half artificial correct trial was added in the case that all trials were incorrect (Macmillan and Creelman, 2004).

### MEG data preprocessing and analysis

The MEG data analysis was conducted in MATLAB using the Fieldtrip toolbox 20170830 (Oostenveld et al., 2011) and wavelet toolbox. Raw MEG data were noise-reduced offline using the time-shifted principle component analysis (de Cheveigné and Simon, 2007) and sensor noise suppression (de Cheveigné and Simon, 2008). Trials were visually inspected, and those with artifacts such as signal jumps and large fluctuations were discarded. An independent component analysis was used to correct for eye blink-, eye movement-, heartbeat-related and system-related artifacts. Twenty trials were included in the analysis for each frozen sound and 30 trials were included in the analysis for distinct sounds of each stimulus type. Each trial was divided into 4 s epochs, with a 1.5 s pre-stimulus period and 2.5 s post-stimulus period. Baseline was corrected for each trial by subtracting out the mean of the whole 4 s trial before further analysis.

To extract time-frequency information, single-trial data in each MEG channel were transformed using functions of the Morlet wavelets embedded in the Fieldtrip toolbox, with a frequency range from 1 to 60 Hz in steps of 1 Hz. To balance spectral and temporal resolution of the time-frequency transformation, from 1 to 20 Hz, window length increased linearly from 2 cycles to 10 cycles, and was then kept constant at 10 cycles above 20 Hz. Phase and power (squared absolute value) were extracted from the wavelet transform output at each time-frequency point.

The ‘inter-trial phase coherence’ (ITPC), a measure of consistency of phase-locked neural activity entrained by stimuli across trials, was calculated on each time-frequency point (details as in Lachaux et al., 1999) for 20 trials of each frozen sound to test neural entrainment on each neural band. ITPC was also calculated for the first 20 distinct sounds of each stimulus type to provide a baseline for each neural band. Robust phase-locking on each band can be potentially affected by the power profiles of MEG signals, since high ITPC values in a given neural band may result from its high power magnitude rather than from high phase coherence across trials. If ITPCs of the distinct sounds do not show an effect of phase coherence in a neural band but ITPCs from 20 trials of each frozen sound manifest such an effect, we could conclude that the robust neural entrainment reflected by ITPC is not confounded by the strength of power of the neural band. As ITPC is not normally distributed, the rationalized arcsine transform was applied on ITPC values before further analyses and statistic tests on ITPC (Studebaker, 1985).

The evoked power response, which reflects phase-locked neural responses, was computed for each frozen sound by applying the time-frequency transform on an averaged temporal response across 20 trials for each frozen sound. Then, the power values were normalized by dividing the mean power value in the baseline range (-.7 ~ -.3 s) and taking logarithms with base 10, and then was converted into values with unit of decibel by multiplying by 10. The evoked power was also calculated for 20 distinct sounds of each stimulus type, which was used as baselines to determine significant power responses for the frozen sounds.

The induced power response was calculated for 20 distinct sounds of each stimulus type to match the analysis conducted for the frozen sounds. The baseline correction was the same as used in calculations of the evoked power. We only chose to compute induced power response on distinct sounds because the induced power response calculated from the frozen sounds contains evoked response components and cannot be fully differentiated from the evoked power response. As distinct sounds of each stimulus type have different temporal structures (modulation phases), the evoked component can be, theoretically, averaged out. Furthermore, we calculated induced power responses without baseline correction to show whether the raw power spectra are different for different stimulus types, which may bias ITPC estimations.

The ITPC and power data were averaged from 0.3 s to 1.8 s post-stimulus onset to minimize the effects of stimulus-evoked onsets and offsets, and within five frequency bands for presenting results of topographies of ITPC: theta (4 - 7 Hz), alpha (8 – 12 Hz), beta1 (13 – 20 Hz), beta2 (21 – 30 Hz) and gamma1 (31 – 45 Hz). As the ITPC analysis showed prominent phase-locking effects for *β2* sounds from 40 – 52 Hz in our later analyses, we further arbitrarily defined the gamma2 band (40 – 52 Hz). We referred to frequencies above 30 Hz as the gamma band, which includes the gamma1 and gamma2 bands.

All calculations were first conducted in each MEG channel and then averaged across 20 selected auditory channels. Statistical analyses of ITPC and power were conducted separately for the frozen sounds and the distinct sounds using repeated-measures ANOVA (rmANOVA) across stimulus types on each frequency point. When multiple comparisons were performed, to control false positive rate, adjusted False Discovery Rate (FDR) was used (Benjamini and Hochberg, 1995; Yekutieli and Benjamini, 1999).

### Single-trial classification

Single-trial classification analysis of frozen sounds for each stimulus type was carried out for each neural band to examine how the auditory system encodes temporal information at different timescales. The procedure was described in detail in Ng et al. (2013), and similar methods were also used in Luo & Poeppel (2012), Herrmann et al. (2013), Cogan et al. (2011) and Teng et al. (2017). For three frozen sounds of each stimulus type, one trial was left out for each sound, and then a template was created by averaging across the remaining 19 trials for this sound (the circular mean is used for phase average). Three templates were created, and the distance between each template and each left-out trial of each frozen sound was computed. The circular distance was applied for phase classification by taking the circular mean over time (0.3 – 1.8 s) and frequencies within each neural band; the l_2_ norm of the linear distance was used for power classification. A left-out trial was given the label of one template if the distance between this trial and the template was the smallest among three templates.

A confusion matrix of classification was constructed by carrying out classification for each trial of each frozen sound of each stimulus type on each auditory channel. Then, classification performance of each neural band on each stimulus type was measured by converting confusion matrices to *d*′: correctly labeling the target frozen sound was counted as a ‘hit’ while labeling the other two frozen sounds as the target frozen sound was counted as a ‘false alarm’; *d*′ was calculated based on hit rates and false alarm rates and averaged across all auditory channels. An index of classification efficiency using phase and power response of different frequency band was indicated by the mean of *d*′ over three frozen sounds of each stimulus type, which was compared to the total *d*′ of the identification task (Macmillan and Creelman, 2004).

### MEG source reconstruction

As high resolution structural T1-weighted MRI scans were only acquired for eight participants, we conducted source reconstruction and localized ITPC and classification efficiency for these participants.

Head shape and head position measurements were taken before the MEG recording session. Both head shape and position were used to co-register individual brain models to each subject’s head using uniform scaling, translation and rotation. The source reconstructions were done by estimating the cortically constrained dynamic statistical parametric mapping (dSPM) of the MEG data. The forward solution (magnetic field estimates at each MEG sensor) was estimated from a source space of 5121 activity points with a boundary-element model (BEM) method. The inverse solution was calculated from the forward solution. Subsequently, we morphed each individual brain to the FreeSurfer average brain (CorTechs Labs Inc., Lajolla, CA) and then averaged ITPC and classification efficiency results across eight participants.

We conducted time-frequency analysis to extract phase series and computed ITPC using MNE-Python in source space with parameters comparable to the time-frequency analysis described above on sensor space (Gramfort et al., 2014). We computed ITPC for each frozen sound using 20 trials and then averaged ITPCs from all three frozen sounds for each stimulus type. We selectively presented source reconstructions of ITPC of each stimulus type in the neural bands that showed significant phase locking effects in the ITPC analysis on the sensor level (*θ* sounds in the theta band, *α* sounds in the alpha band, *β1* sounds in the beta1 band, *β2* sounds in the beta2 and gamma2 bands, *γ* sounds in the gamma1 band). For single-trial classification, phase series were first exported from Python to Matlab and classification efficiency for each stimulus type was calculated using the same procedures as in *single-trial classification*.

### Stimulus reconstruction

To investigate how acoustic information of different timescales is faithfully encoded by each neural band, we reconstructed cochleograms of different stimulus types from each neural band. The underlying hypothesis is that, if a neural band encodes acoustic dynamics characteristic of a stimulus type, this neural band can be used to reconstruct the cochleograms of this stimulus type with high accuracy while other neural bands that do not encode acoustic dynamics can only aid in reconstructing the stimulus type to a limited extent.

The method used here is to map between cochleograms of stimuli and the MEG signals. A temporal response function (TRF) was derived from the cochleograms of stimuli (*S(t)* with subscript *c* indicating cochlear band number) and their corresponding MEG signals (*R(t)* with subscript *b* indicating neural band) through ridge regression with a parameter (lambda) to control for overfitting (superscript *t* indicating transpose operation):

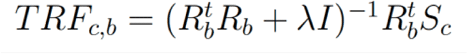

Cochleograms were reconstructed from TRF models as:

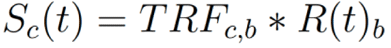

The reconstruction process included two stages: a training stage and a testing stage (illustrated in Fig. 5*A*). At the training stage, we used 30 distinct sounds of each stimulus type and their corresponding MEG recordings as a training set to derive TRFs and then used 10 trials from each of three frozen sounds as a validation set to determine the optimal lambda which gave the highest reconstruction performance. At the testing stage, we applied the derived TRFs and lambda values to the remaining 10 trials of each of three frozen sounds and reconstructed cochleograms of the frozen sounds. Each reconstructed cochleogram was compared with its original cochleogram, and then model performance was measured by computing Pearson correlation (r) between them. Reconstruction performances from the test set for three frozen sounds of one stimulus type were then averaged. We used the distinct sounds as training samples instead of the frozen sounds because three frozen sounds for each stimulus type represent limited variations of acoustic dynamics while 30 distinct sounds cover a wide range of variations of dynamics for each stimulus type – each distinct sound is different from the other distinct sounds. Therefore, training using the distinct sounds is not biased towards a specific sample. TRFs were calculated using the Multivariate Temporal Response Function (mTRF) Toolbox (Crosse et al., 2016).

**Figure 5.**
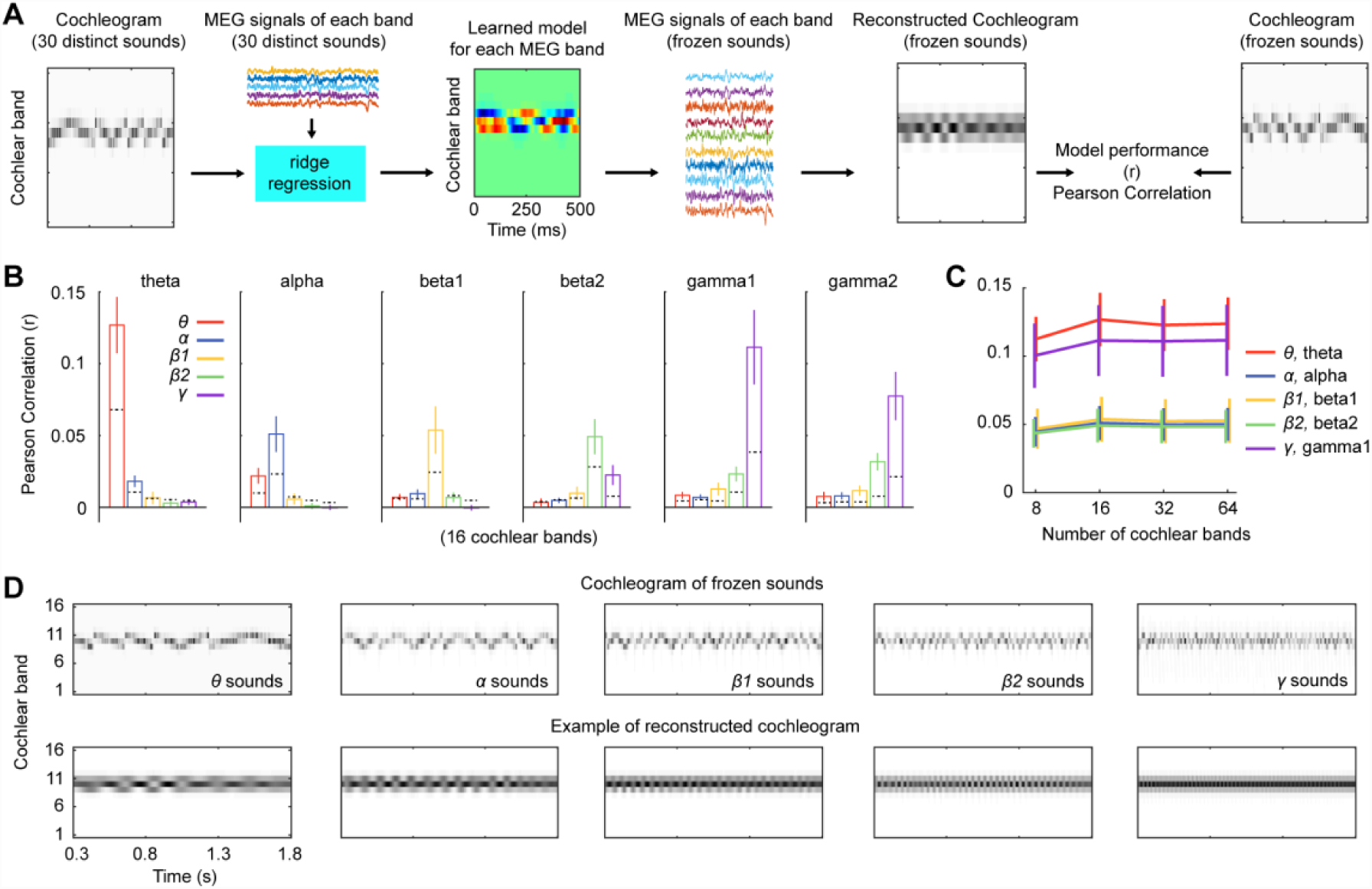
Stimulus reconstruction. ***A***, illustration of procedures of stimulus reconstruction. Thirty distinct sounds of a stimulus type were used to train a TRF model which was then validated using 10 trials of frozen sounds of this stimulus type to estimate an optimal lambda (see Method for details). The TRF model was tested using the remaining 10 trials of frozen sounds and the model performance was quantified as Pearson correlation between the reconstructed cochleograms of frozen sounds and their original cochleograms. ***B***, stimulus reconstruction using 16 cochlear bands within each neural band for all the stimulus types. The color scheme codes for different stimulus types and the dashed line within each bar represents the permutation threshold of alpha level of 0.01. ***C***, stimulus reconstruction using different numbers of cochlear bands on each stimulus type within its corresponding neural band. The color scheme codes for each stimulus type within its corresponding neural band. The *θ* and *γ* sounds can be reconstructed from the respective neural bands with comparably high performance compared with the other stimulus types. ***D***, examples of reconstructed cochleograms from one subject. From the reconstructed cochleograms, it can be seen that modulation patterns of different stimulus types are preserved.

The cochleograms for reconstruction were generated using 8, 16, 32, and 64 bands, separately, ranging from 50 Hz to 4000 Hz (methods described above). This frequency range includes most of the spectral energy of our stimuli centered between 1000 Hz and 1500 Hz. MEG signals were decomposed into theta, alpha, beta1, beta2, gamma1 and gamma2 bands using a two-pass Butterworth filter with an order of 2 embedded in the Fieldtrip toolbox. Each cochlear band was reconstructed individually from each neural band. The model performance was calculated for each cochlear band and then averaged across all cochlear bands.

One concern with regard to the procedure of stimulus reconstruction here is that the TRF model trained on the basis of a neural band may yield a good stimulus reconstruction merely because the frequency of that neural band overlaps with the modulation rate of the stimuli. To control for this confound, we conducted a permutation test. All the procedures of stimulus reconstruction remained the same in the permutation test, but the pairings in the training set between the distinct sounds and their corresponding MEG responses were shuffled, yielding a new set in which each distinct sound was paired with an MEG response to a different distinct sound. We conducted this permutation test 500 times and derived a permutation threshold with a one-sided alpha level of 0.01 for each stimulus type and each neural band.

## Results

### Behavioral performance

Participants discriminated between *θ*, *α*, *β1*, *β2*, and *γ* sounds in the match-to-sample task, and the results show that the discriminability between different modulation rates increases as the difference between two modulation rates becomes larger (Fig. 1*C*, upper panel). The discriminability between adjacent modulation rates is best at the low frequency range and decreases as modulation rates increases, which can be seen from the perceptual distances between stimulus types (Fig. 1*C*, lower panel). For example, although the difference of modulation rates between *θ* and *α* sounds is smaller than between *β2* and *γ* sounds*, θ* and *α* sounds were better differentiated. We selected d-prime values of four pairs of adjacent modulations (*θ* vs. *α*; *α* vs. *β1*; *β1* vs. *β2*; *β2* vs. *γ*) and conducted a one-way rmANOVA with modulation rate difference between two stimulus types as the factor. We found a significant main effect (*F*(3,42) = 5.11, *p* = 0.040, η_p_^2^ = .267).

### Phase coherence and power responses for frozen and distinct sounds

The ITPC values for the frozen sounds in response to each stimulus type from 2 to 60 Hz are plotted in Fig. 2*A*. The results of ITPC show robust neural entrainment for all the five stimulus types in their corresponding neural bands, with *θ* and *γ* sounds in the corresponding bands showing the most prominent phase-locking effects. The much reduced phase coherence for *α*, *β1*, and *β2* sounds in the respective neural bands is inconsistent with the simplest hypothesis that acoustic dynamics across all temporal regimes are faithfully tracked in the auditory cortex. Interestingly, an effect of phase tracking is also observed for *β2* sounds in the gamma2 band (40 – 52 Hz), which we examined further in subsequent analyses. The phase-locking patterns across temporal regimes are mirrored in topographies of ITPC (Fig. 2*B*), which manifest clear auditory response patterns for *θ* sounds in the theta band, *γ* sounds in the gamma1 band, and *β2* sounds in the gamma2 band. The results of evoked power are consistent with ITPC and show robust power responses across frequency and time for *θ* and *γ* sounds in the theta band and gamma band, respectively (Fig. 2*D*). In contrast, ITPC and power results for distinct sounds do not show any effects in different neural bands, and, importantly, the power spectra without baseline correction and the induced power spectra are comparable across all stimulus types (Fig. 2*E*).

**Figure 2.**
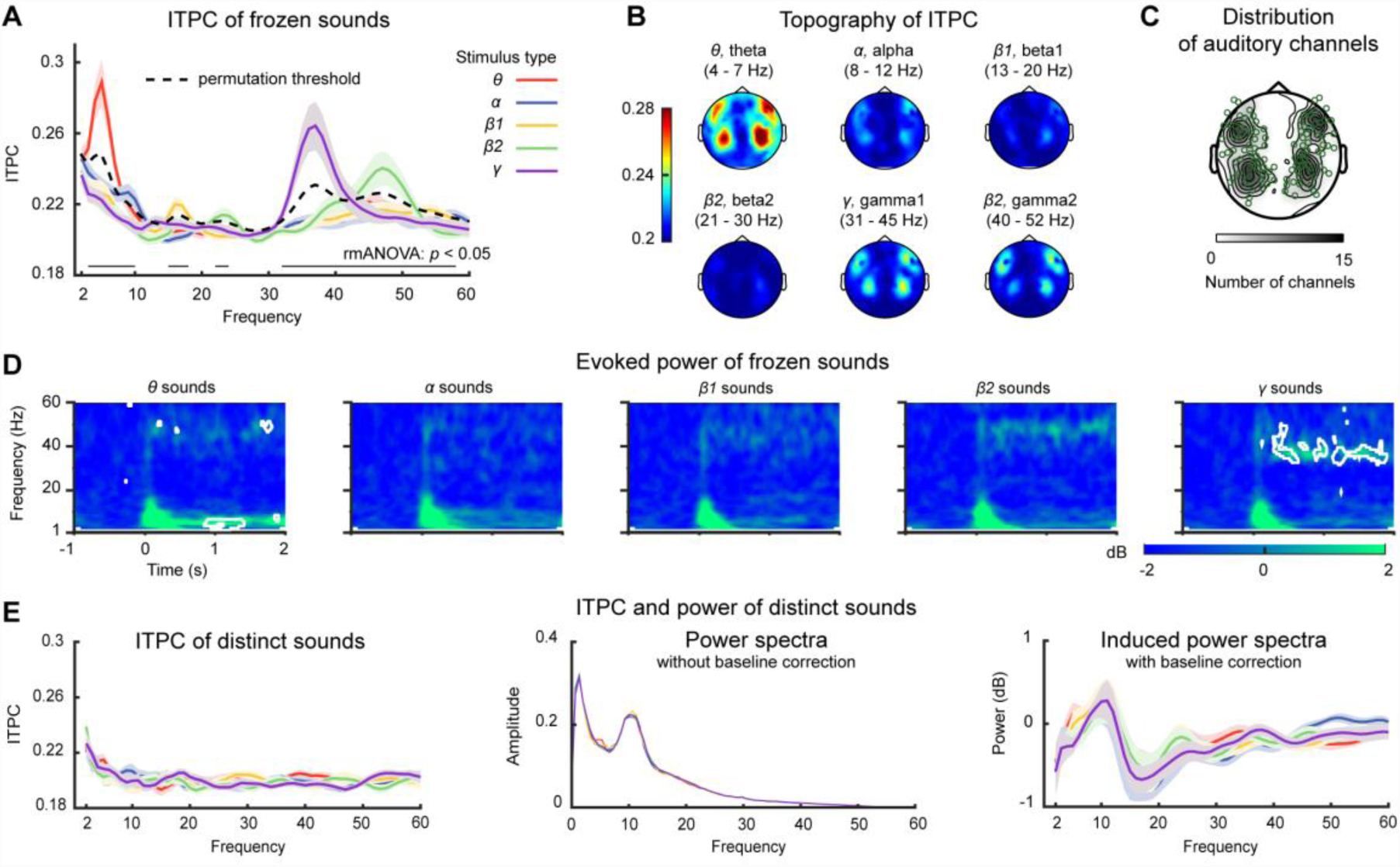
ITPC and power results. ***A***, Spectra of ITPC for the frozen sounds of five stimulus types. The color scheme codes for *θ*, *α*, *β1*, *β2*, and *γ* sounds, respectively. The shaded areas represent ±1 standard error of the mean across participants. The dashed line represents the permutation threshold with a one-sided alpha level of 0.01 (see details in the main text). ITPC values of the stimulus type above the permutation threshold indicate that the stimulus type evoked significantly higher ITPC values than the other stimulus types. The thin solid line above the x-axis indicates frequencies where the main effect of the stimulus type, tested by rmANOVA, is significant (*p* < 0.05, FDR corrected). The results show significant neural entrainment for all the five stimulus types in their corresponding neural bands, but with the theta band for *θ* sounds and the gamma band for *γ* sounds showing the most prominent phase-locking effects. ***B***, Topographies of ITPC for each stimulus type. Patterns of typical auditory responses can be seen for *θ* sounds in the theta band, *γ* sounds in the gamma1 band, and *β2* sounds in the gamma2 band. ***C***, A layout of MEG channels that are selected based on the peak of M100 response. Twenty channels are selected for each subject (10 in each hemisphere). The channels selected for analysis are indicated by circles. The contours indicate the extent of overlap of selected auditory channels across participants. ***D***, Evoked power for the frozen sounds for each stimulus type. From left to right, each panel shows evoked power responses for *θ*, *α*, *β1*, *β2*, and *γ* sounds, respectively. The white contours indicate spectral-temporal tiles where the evoked power responses of the frozen sounds are significantly larger than the evoked power computed using twenty distinct sounds (*p* < 0.05, FDR corrected). ***E***, ITPC and power results for the distinct sounds. Left panel shows spectra of ITPC for the distinct sounds for each stimulus type. Middle panel shows power spectra without baseline correction. Right panel shows induced power spectra (with baseline correction). It can be seen that the power responses of the distinct sounds do not differ between the stimulus types and hence do not contribute to the ITPC results in different neural bands.

#### Phase coherence

We first averaged ITPC values for each stimulus type across five pre-defined neural bands - theta (4 - 7 Hz), alpha (8 – 12 Hz), beta1 (13 – 20 Hz), beta2 (21 – 30 Hz), gamma1 (31 – 45 Hz), and one neural band defined post hoc - gamma2 (40 – 52 Hz). We conducted a Stimulus-type × Hemisphere × Neural band three-way rmANOVA on ITPC. This revealed main effects of Stimulus type (*F*(4,56) = 9.06, *p* < 0.001, η_p_^2^ = .392), Neural band (*F*(5,70) = 20.92, *p* < 0.001 η_p_^2^ = .599) and Hemisphere (*F*(1,14) = 8.34, *p* = 0.012, η_p_^2^ = .373). The interaction between Stimulus-type and Neural band is significant (*F*(20,280) = 20.70, *p* < 0.001, η_p_^2^ = .597). Although the main effect of Hemisphere is significant with ITPC values moderately higher in the right hemisphere than the left hemisphere (left: 0.2131 ± 0.0028; right: 0.2166 ± 0.0028), no significant interaction effects were found between Hemisphere and Neural band (*F*(5,70) = 1.55, *p* = 0.186, η_p_^2^ = .100) and between Hemisphere and Stimulus type (*F*(4,56) = 1.31, *p* = 0.279, η_p_^2^ = .085). Therefore, in the analyses to follow, we combined all the selected auditory channels (Fig. 2*C*) from both hemispheres.

To investigate the differences of neural entrainment between different stimuli types in different neural bands, we conducted a Stimulus-type one-way rmANOVA on each frequency point of ITPC spectra from 2 to 60 Hz averaged across all auditory channels. This revealed a main effect of Stimulus-type (*p* < 0.01) in the frequencies of 2 – 10 Hz, 15 – 18 Hz, 22 – 24 Hz, and 32 – 58 Hz (Fig. 2*A*) (FDR correction was applied).

To further examine which stimulus type yielded the highest ITPC values in the significant frequency ranges shown above (2 – 10 Hz, 15 – 18 Hz, 22 – 24 Hz, and 32 – 58 Hz), we conducted a permutation test, in which we shuffled the labels of the frozen sounds for ITPC values of each subject and repeated the procedure to derive a new ITPC value for each stimulus type. The shuffled ITPC values were then averaged across 15 participants to derive a group mean. Through this shuffling process, the ITPC values for frozen sounds are randomly grouped into each stimulus type and the shuffled ITPC values can be used as baselines for the true ITPC values of each stimulus type. We repeated the shuffling procedure 500 times and derived a one-sided alpha level of 0.01 as a threshold of the group mean of ITPC for each stimulus type. As the shuffling procedure treated each stimulus type without bias, we then averaged the thresholds of the five stimulus types as a single threshold for ITPC for all the stimulus types. If the ITPC of one stimulus type is above this threshold, we conclude that the ITPC of this stimulus type is significantly larger than the other stimulus types. We found that, within the frequency ranges that show significant main effects of Stimulus-type, ITPC of *θ* sounds is significantly above the threshold from 3 to 8 Hz, as well as *α* sounds from 8 to 10 Hz and from 57 to 58 Hz, *β1* sounds from 15 to 18 Hz and from 54 to 56 Hz, *β2* sounds from 22 to 24 Hz and from 43 to 51 Hz, *γ* sounds from 32 to 42 Hz. In summary, significant effects of phase coherence are found for all stimulus types within their respective neural bands and, surprisingly, in the gamma band range for *α, β1,* and *β2* sounds.

Since high ITPC values can be simply caused by high power in a neural band but not by phase coherence across trials, to provide a baseline of ITPC we calculated ITPC using the same procedures on 20 distinct sounds for each stimulus type and performed a Stimulus-type one-way rmANOVA on each frequency point of ITPC spectra from 2 to 60 Hz (Fig. 2*E*, left panel). No significant main effect of Stimulus-type was found after FDR correction (*p* > 0.05).

#### Evoked power response

We conducted paired t-tests to compare evoked power between the frozen sounds and the distinct sounds. The paired t-tests were conducted on each time-frequency point from −0.5 s to 1.9 s and from 2 Hz to 60 Hz and FDR correction was applied. We cut off the power responses from 1.9 s to 2 s because at low frequencies (e.g. 2 Hz) large temporal windows used in wavelet analysis did not give a valid estimate of power responses due to the epoch size of each trial. In Fig. 2*D*, we used the white contours to indicate the time-frequency points where significant differences of evoked power between the frozen sounds and the distinct sounds were found. Although salient power responses can be observed in the theta and gamma bands for all stimulus types, significant power responses (*p* < 0.05) entrained by the frozen sounds were mainly found in the theta band for *θ* sounds and in the gamma1 band for *γ* sounds.

#### Induced power response for the distinct sounds

We calculated induced power with and without baseline correction for the distinct sounds of each stimulus type (Fig. 2*E*, middle and right panels). The rationale for this analysis was that the components of evoked power could be conceivably averaged out, as the distinct sounds have different modulation phases from each other. Therefore, we could examine, without the influence of time-locked components, how different stimulus types yield different power responses. We performed a Stimulus-type one-way rmANOVA on each frequency point of power spectra and found no significant effects of Stimulus-type after FDR correction (*p* > 0.05). This result demonstrates that induced power responses are comparable across the five stimulus types and do not contribute to estimation of neural entrainment.

### Classification using phase and power for the frozen sounds

Classification efficiency of each neural band for each stimulus type was calculated using the MEG signals in both the phase and power domains. If the MEG response from a particular neural band can differentiate three frozen sounds of one stimulus type, this would suggest that this neural band encodes detailed temporal information of this stimulus type (modulation phase of each sound). The results of classification efficiency, surprisingly, demonstrates that the phase information in both the theta and gamma bands reliably differentiated the frozen sounds of all the five stimulus types, while the alpha and beta bands show only moderate classification performance (Fig. 3*A*). Further analyses show that the frozen sounds of all the stimulus types can be differentiated with comparable accuracy using phase information of all the frequency bands together (2 – 60 Hz) (Fig. 3*C*, left panel) and the classification performance is contributed to primarily by the theta and gamma bands (Fig. 3*C*, right panel). Together, the results of ITPC, evoked power, and classification show that acoustic dynamics of all timescales used in the present study are encoded mainly by the theta and gamma bands.

**Figure 3.**
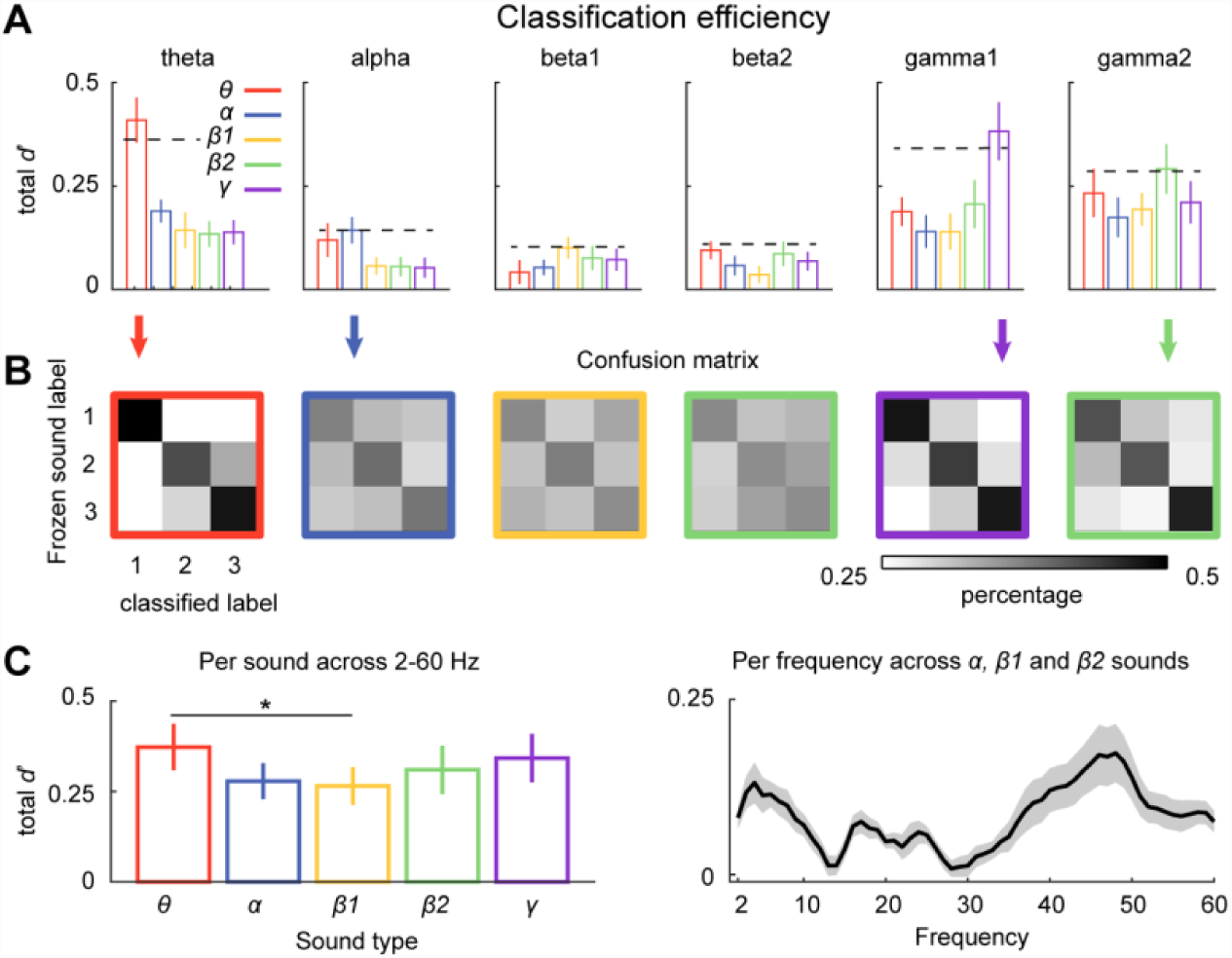
Classification results using phase information of the MEG response. ***A***, classification efficiency for each stimulus type within different neural bands. The color scheme is the same as in Fig. 2. From left to right, each plot shows classification efficiency of each neural band. The dashed line is the permutation threshold (see details in the main text). The results show robust classification performance in the theta and gamma bands for all the stimulus types, while moderate performance is shown in the alpha, beta1 and beta2 bands. ***B***, group-averaged confusion matrices for each stimulus type in its corresponding bands. The color of the contours of confusion matrices codes for stimulus types. The neural bands represented by each of the confusion matrices are indicated by the arrows and align with the neural bands of ***A***. ***C***, classification efficiencies of full bands and per frequency. Left panel shows classification efficiency for each stimulus type computed using full bands of phase information in the MEG signals and demonstrates that the frozen sounds of all stimulus types are comparably classified, although much reduced neural entrainment is observed for *α*, *β1*, and *β2* sounds (Fig. 2). Right panel shows classification efficiency of *α*, *β1*, and *β2* sounds per frequency. Two prominent peaks can be clearly seen in the theta and gamma ranges. This suggests that the major contribution to classification performance comes from theta and gamma bands.

We first test whether phase and power information of each neural band contributes to differentiation of frozen sounds of each stimulus type by conducting a one-sample t-test against zero for classification efficiency of each neural band and each stimulus type. After FDR correction was applied, we found that none of classification efficiencies calculated using power information were significantly above zero for any stimulus types and any neural bands (*p* > 0.05). In contrast, classification efficiencies calculated using phase information were significantly above zero for all the stimulus types and all the neural bands (*p* < 0.05) except for *θ* sounds in the beta1 band (*t*(1,14) = 1.46, *p* = 0.164), *β1* sounds in the beta2 band (*t*(1,14) = 1.76, *p* = 0.100) and *γ* sounds in the alpha band (*t*(1,14) = 2.17, *p* = 0.051). Therefore, in the following analyses, we only investigated classification efficiencies calculated using phase information.

To determine for each neural band which stimulus type is preferably encoded and therefore has the highest classification efficiency, we did a permutation test similar to the one that we conducted for testing ITPC values. We shuffled the labels of 15 frozen sounds within each neural band and assigned three frozen sounds randomly chosen from the 15 frozen sounds to each stimulus type. By doing this for each subject, we created a new set of classification efficiencies for each stimulus type; the group mean for each stimulus type was then computed. We repeated this procedure 500 times and derived a one-sided alpha level of 0.01 for each stimulus type. As the shuffling procedure was unbiased to any stimulus types, we averaged the thresholds across the five stimulus types and created a single threshold for each neural band. If classification efficiency for one stimulus within one neural band is above the derived threshold, we conclude that this stimulus type is preferentially encoded and significantly better classified than the other stimulus types. We found that *θ* sounds in the theta band, *α* sounds in the alpha band, *γ* sounds in the gamma1 band, and *β*2 sounds in the gamma2 band have classification efficiencies above the permutation thresholds.

One observation from Fig. 3*A* is that the classification efficiencies are much higher for all the stimulus types in the theta and gamma bands compared with the other neural bands. We suspect that the frozen sounds of all the stimulus types are probably classified with comparable efficiencies and are mainly encoded by the theta and gamma bands. To test this, we first computed classification efficiency for each stimulus type using frequencies from 2 to 60 Hz, which included all the neural bands, and conducted paired t-tests between each pair of stimulus types. After FDR correction was applied, we found a significant difference of classification efficiency between *θ* sounds and *β*1 sounds (t(1,14) = 3.96, *p* = 0.014), but not between other stimulus types (*p* > 0.05) (Fig. 3*C*, left panel). This result proves that, although small differences of classification efficiency exist between stimulus types, acoustic dynamics of all the stimulus types are comparably encoded by the neural signals recorded by MEG. This result raises a question: what neural bands are encoding acoustic dynamics of *α, β1,* and *β2* sounds if neural entrainments for these three stimulus types are much reduced (Fig. 2*A*)? We then conducted classification analyses using phase information of each frequency and averaged classification efficiencies across *α, β1,* and *β2* sounds to show a spectrum of classification efficiency (Fig. 3*C*, right panel), which echoes Fig. 3*A* and shows two prominent peaks of classification efficiency within the theta and gamma band ranges.

The classification results demonstrate that acoustic dynamics of all temporal ranges are primarily encoded by the theta and gamma bands, to a comparable degree. The fact that the theta band encodes temporal information of different timescales is probably a result of a ‘chunking’ or segmentation process - the auditory system actively chunks sounds into segments of around 150–300 ms, roughly a cycle of the theta band, for grouping acoustic information (Ghitza and Greenberg, 2009; Ghitza, 2012; Teng et al., 2018). Although different stimulus types have acoustic dynamics on different timescales, this chunking process actively groups acoustic information into chunks in the approximate time window of a theta period. The theta band signals reflect this chunking process and therefore can be used to classify all stimulus types (Teng et al., 2018). On the other hand, each gamma cycle integrates fine-grained acoustic information on a local scale (e.g. ~ 30 ms), for example, transient segment onsets in the stimuli. Therefore, the gamma band reflects encoding temporal information of each stimulus type on a local scale, in comparison with the theta range, and can also be used to classify all the stimulus types (Poeppel, 2003).

### Source localization of ITPC and classification efficiency

We conducted source reconstruction of MEG signals for 8 participants with available MRIs and projected the results of ITPC and classification efficiency to source space. Fig. 4 shows source plots of ITPC and classification efficiency averaged across 8 participants. High ITPC values for all the stimulus types are centered around auditory cortical areas, and considerable neural entrainment can be observed for all the stimulus types with *θ*, *α* and *γ* sounds showing higher ITPC values than *β1* and *β2* sounds (Fig. 4*A*). The results of classification efficiency (Fig. 4*B*), compared with ITPC results, show a different pattern – robust classification performance can only be seen for *θ* sounds in the theta band and *γ* sounds in the gamma1 band but not for *α*, *β1* and *β2* sounds.

**Figure 4.**
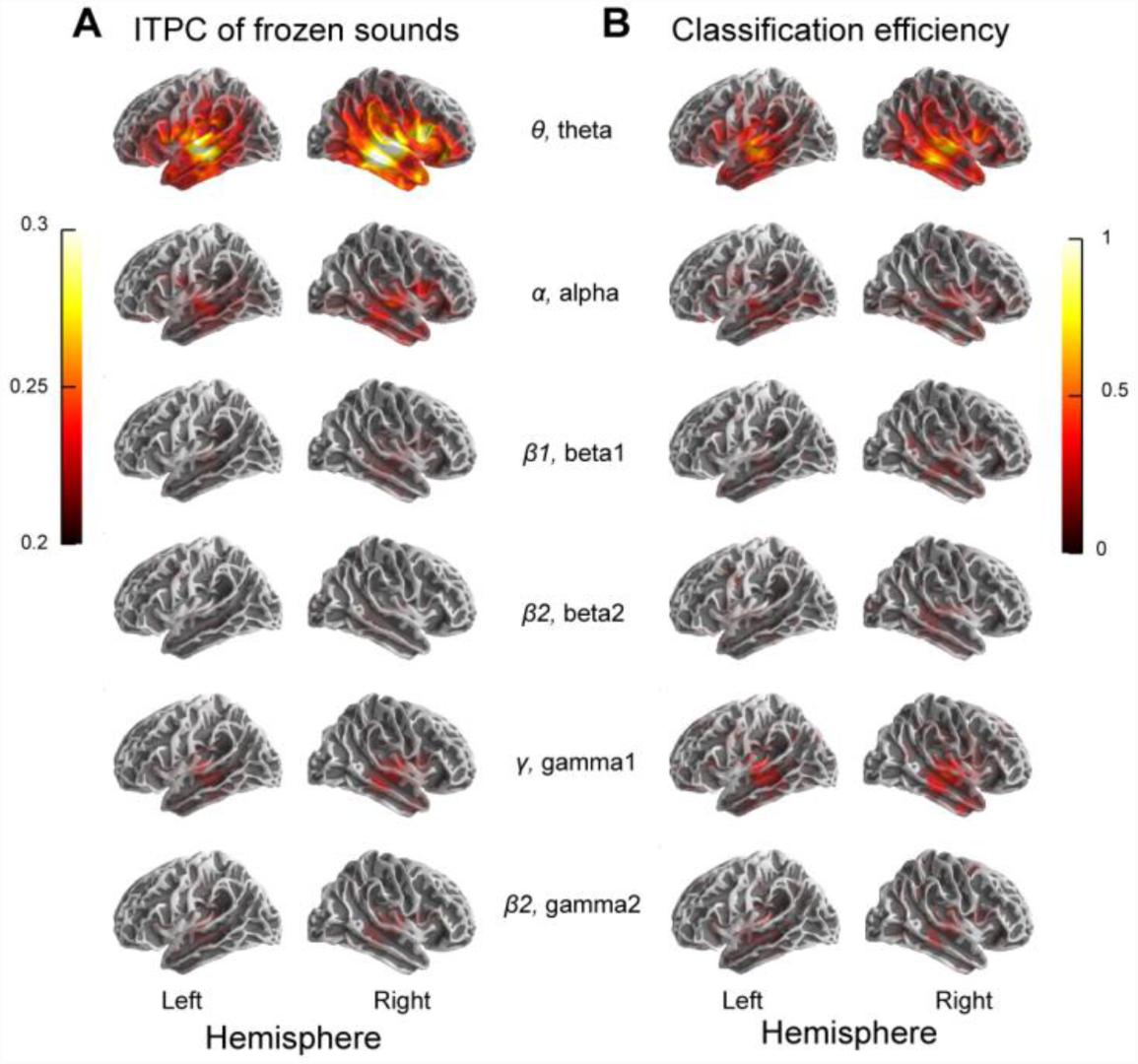
Source localization of ITPC and classification efficiency from 8 participants. ***A***, source plots of ITPC for the frozen sounds. High ITPC values are centered around auditory cortical areas, which demonstrates that robust neural entrainment to different stimulus types originates from similar auditory cortical areas. ***B***, source plots of classification efficiency for the frozen sounds. Considerable classification performance can be seen for *θ* sounds in the theta band and *γ* sounds in the gamma1 band around auditory cortical areas. Moderate classification performance can also be seen for *α, β1,* and *β2* sounds, but the magnitude is much reduced compared with *θ* and *γ* sounds. The legends between ***A*** and ***B*** indicate the stimulus type and the neural band. For instance, ‘*θ*, theta’ indicates that the ITPC values and the classification efficiency on that row represent the results of *θ* sounds in the theta band.

### Stimulus reconstruction

The classification analysis showed that the theta and gamma bands aided in classifying the frozen sounds of different stimulus types (Fig. 3), but this analysis was conducted only on the neural signals, without directly relating the neural signals to the acoustic details of stimuli. It is still unclear whether the theta and gamma bands directly encode acoustic details of all the stimulus types or only code temporal information reflected in the modulation rates of the stimuli. Using stimulus reconstruction from the MEG signals, we next examine how neural signals of each band specifically encode acoustic dynamics of each stimulus type.

The procedures of stimulus reconstruction are shown in Fig. 5*A*. We first reconstructed cochleograms of 16 bands for each stimulus type, to investigate how well each stimulus type can be decoded from different bands of the MEG signals (Fig. 5*B*). The results show that each stimulus type can be reconstructed sufficiently by its corresponding neural band. We further varied the number of cochlear bands and found that the number of cochlear bands modulates stimulus reconstruction, and that *θ* and *γ* sounds are reconstructed with comparably high accuracy from their corresponding neural bands (Fig. 5*C*). Fig. 5*D* shows examples of reconstructed cochleograms for each stimulus type from one subject.

The performance of stimulus reconstruction using 16 cochlear bands was compared using the threshold of alpha level, 0.01, derived from permutation (see Method for details). The results (Fig. 5*B*) show significant reconstruction performance for *θ*, *α*, and *β1* sounds in the theta band, *θ* and *α* sounds in the alpha band, *θ*, *α* and *β1* sounds in the beta1 band, *β1, β2* and *γ* sounds in the beta2 band, and all the stimulus types in the gamma band (gamma1 and gamma2). The highest reconstruction performance was observed for each stimulus type in its corresponding band. We then focused on the reconstruction performance of each stimulus type from its corresponding band using different numbers of cochlear bands. We conducted a Stimulus-type × Cochlear band two-way rmANOVA on reconstruction performance. As reconstruction performance was measured by Pearson correlation, a z-transform on the correlation coefficient was applied before the rmANOVA. We found significant main effects of Stimulus-type (*F*(4,56) = 9.86, *p* < 0.001, η_p_^2^ = .413) and Cochlear band (*F*(3,42) = 18.23, *p* < 0.001, η_p_^2^ = .566), as well as a significant interaction effect (*F*(32,168) = 3.37, *p* < 0.001, η_p_^2^ = .194). Post-hoc comparisons using paired t-tests with FDR correction on the main effect of Stimulus-type show that reconstruction performance of *θ* and *γ* sounds are significantly larger than other stimulus types (*p* < 0.05) but not different from each other (*t*(1,14) = −0.77, *p* = 0.646). The linear trend of Cochlear band is significant (*F*(1,14) = 17.90, *p* < 0.001, η_p_^2^ = .561), suggesting that the decoding performance increases with the number of cochlear bands used in the stimulus reconstruction.

The results of stimulus reconstruction demonstrate that the theta and gamma bands specifically encode the acoustic details of *θ* and *γ* sounds, respectively. The reconstruction performance for *α*, *β1* and *β2* sounds is significant, but is much lower than for *θ* and *γ* sounds. Although the theta and gamma bands show an effect for all the stimulus types in the *classification* analysis (Fig. 3), such a generality of temporal coding is not observed in the *stimulus reconstruction*. Therefore, it is plausible to conclude that the theta and gamma bands code temporal information at all timescales represented by the modulation rates of the stimuli, but preferably extract acoustic features on two timescales: ~ 30 ms and ~ 200 ms.

## Discussion

We show that acoustic dynamics are reliably tracked in the theta and gamma bands but not in the alpha and beta bands (Fig. 2), consistent with earlier findings (Luo and Poeppel, 2012; Wang et al., 2012; Teng et al., 2017). Classification analyses showed that the neural theta and gamma bands contribute to the differentiation of sounds for all stimulus types - and with comparable temporal coding capacity (Fig. 3). Source localization results of ITPC and classification efficiency revealed a similar origin of both the neural theta and gamma bands around auditory cortical areas (Fig. 4). Stimulus reconstruction further supported that acoustic dynamics are faithfully encoded by the theta and gamma bands with comparable precision, but only modestly by the alpha and beta bands (Fig. 5). Following previous work, the results demonstrate that the theta and gamma bands code temporal information of all timescales in general and especially extract acoustic features on the timescales of ~ 30 ms and ~ 200 ms. This provides convincing evidence for the hypothesis that the human auditory system primarily analyzes information in two distinct temporal regimes that carry perceptually relevant information (Poeppel, 2003).

Previous studies on auditory temporal processing typically focused on one temporal regime, mostly in the low frequency range (< 10 Hz). Using various acoustic stimuli, a majority of studies on neural entrainment in auditory cortices suggest a high temporal coding precision primarily in the low frequency range and argue that the temporal coding precision of the auditory system decreases with increased modulation rates (Luo and Poeppel, 2007; Lakatos et al., 2008; Kerlin et al., 2010; Besle et al., 2011; Cogan and Poeppel, 2011; Ding and Simon, 2012; Henry and Obleser, 2012; Kayser et al., 2012; Ng et al., 2012; Wang et al., 2012; Ding and Simon, 2013; Herrmann et al., 2013; Lakatos et al., 2013; Peelle et al., 2013; Doelling et al., 2014; Henry et al., 2014; Kayser et al., 2015; Riecke et al., 2015; Zoefel and VanRullen, 2015). However, several other studies provide an alternative view. A study using amplitude modulation created by binaural beats showed strong entrainment in both the theta and gamma bands (Ross et al., 2014). Recordings in the primary auditory cortex of monkeys also show a phase-locked response using amplitude modulation at 30 Hz (Brosch et al., 2002; Johnson et al., 2012). Gamma band entrainment is also found to contribute to speech separation in multiple speaker environments (Kerlin et al., 2010). Seen in conjunction with our own previous work (Luo and Poeppel, 2012; Teng et al., 2017), the current findings argue for two concurrent temporal channels for auditory processing, with comparable temporal processing capacity. The auditory system tunes to acoustic information at the two discrete timescales with equal temporal precision, which leads to a temporal multiplexing of sensory information (Panzeri et al., 2010; Gross et al., 2013). This may facilitate the efficient extraction of perceptual information at different timescales in speech, such as phonemic scale information and syllabic scale information (Rosen, 1992).

To further examine the argument arising from previous work (Teng et al., 2017), we hypothesize that the marked reduction of temporal coding in the alpha and beta bands suggests that these two bands play a different processing role in the cortical auditory system. It has been well established that, in the auditory system, the computations implied by the neural alpha band may be related to attention, memory load, listening effort, or functional inhibition (Weisz et al., 2011; Obleser and Weisz, 2012; Obleser et al., 2012; Strauß et al., 2014; Wöstmann et al., 2015; Wilsch and Obleser, 2016). Such observations have also been shown in the visual and somatosensory systems (van Dijk et al., 2008; Haegens et al., 2011). Beta band neural signals are argued to play a role in predictive coding (Arnal and Giraud, 2012; Arnal et al., 2015). Therefore, neural coding schemes in the auditory system may be organized based on timescales to optimize sensory input selection (Buzsáki, 2004), with the theta and gamma bands primarily responsible for temporal coding of acoustic information.

The source localizations of ITPC and classification efficiency demonstrate that the neural theta and gamma bands originate from similar auditory cortical areas, which suggests that the two temporal channels coexist in the cortical auditory system. Although we also found activation for alpha and beta bands around similar cortical areas, the temporal coding precision measured by the classification efficiency is sharply reduced in comparison with the theta and gamma bands. Specifically, in the alpha band the results from 8 participants show moderate magnitude of ITPC but much reduced classification efficiency (Fig. 4). This suggests that the reduced temporal coding capacity in alpha and beta bands revealed by our analyses is not because MEG fails to record neural activity in the alpha and beta bands, but because the preferred temporal coding in audition is confined to the theta and gamma ranges.

Our finding of robust temporal coding within the theta and gamma ranges is consistent with previous behavioral studies and has fundamental implications. Two perceptual time constants are often found in behavioral studies (Green, 1985): experiments on temporal integration converge on a time constant of hundreds of milliseconds (Plomp and Bouman, 1959; Green, 1960; Zwislocki, 1960; Jeffress, 1964; Green and Swets, 1966; Jeffress, 1968; Zwislocki, 1969), while studies examining the high temporal resolution of the auditory system show a time constant of less than 30 ms (Viemeister, 1979; Forrest, 1987; Moore, 1988). One recent behavioral study also converges with the present neurophysiological results and demonstrates that the auditory system works concurrently on a short timescale (~30 ms) to extract fine-grained acoustic temporal detail and on a longer timescale to process global acoustic patterns (> 200 ms) (Teng et al., 2016). However, the behavioral task in this study, designed to reveal listeners’ discriminability between different modulation rates, did not yield results in line with our neurophysiological findings. The behavioral results here show a pattern of a low-pass filter shape – listeners’ performance is highest in the low frequency range and decreases with increased modulation rates, which is consistent with the temporal modulation transfer function found in modulation detection paradigms (Dau et al., 1997). The discrepancy between the current behavioral results of modulation discrimination and our neurophysiological results reveals the complicated nature of auditory temporal processing. As the tasks of detection and differentiation of temporal modulations do not require listeners to decipher information embedded in each modulation cycle, the behavioral results of those tasks probably cannot reflect how the auditory system processes acoustic information on each timescale. Our neurophysiological results invite new behavioral paradigms that target auditory processing on timescales between ~ 30 ms and ~ 200 ms.

Natural sounds contain information at multiple scales and, in order to efficiently sample perceptual information, the auditory system chunks continuous sounds using temporal windows of specific sizes instead of processing acoustic information in a continuous manner (Ghitza and Greenberg, 2009; Giraud and Poeppel, 2012). Selective representation of acoustic information using timescales of ~ 30 ms and ~ 200 ms may align with efficient encoding – the auditory system preferably extracts acoustic features of the timescales essential to natural sounds (Lewicki, 2002; Smith and Lewicki, 2006). Such a processing scheme is in line with findings in the visual modality (VanRullen and Koch, 2003; VanRullen, 2006; Blais et al., 2013), for which preferred encoding on ecologically important features is well demonstrated (Olshausen and Field, 2004). One model of auditory processing proposes that - although a very high resolution is represented in sub-cortical areas - on the cortical level of the auditory system there are two main temporal windows used for processing perceptually relevant information: one centered around 200 ms and the other around 30 ms (Poeppel, 2003; Giraud and Poeppel, 2012). Our results on the theta and gamma bands argue for a segregation of function in the auditory system between low and high processing rates – perhaps optimized for sensory sampling – by an intermediate rate perhaps optimized for allocating attentional and memory resources and functionally inhibiting task- or stimulus-irrelevant actions.

## Acknowledgements

We thank Jeff Walker and Jess Rowland for their technical support. This study was supported by NIH 5R01DC005660 to DP and the Max-Planck-Society. We thank Nina Kazanina, Sean Lee, and Ava Kiai for helpful discussions and comments.

**Author contributions** X.T and D.P designed the study. X.T performed the experiment, analyzed the data, and wrote the first draft of the paper; X.T and D.P wrote the paper.

